# Consistency of non-cognitive skills and their relation to educational outcomes in a UK cohort

**DOI:** 10.1101/470682

**Authors:** Tim T Morris, George Davey Smith, Gerard van Den Berg, Neil M Davies

**Author notes:** Corresponding author: Tim T Morris, MRC Integrative Epidemiology Unit, University of Bristol, Oakfield House, Oakfield Grove, BS8 2BN, UK.

## Abstract

Non-cognitive skills have previously been associated with a range of health and socioeconomic outcomes, though there has been considerable heterogeneity in published research. Many studies have used cross sectional data and therefore the longitudinal consistency of measures designed to capture non-cognitive skills is poorly understood. Using data from a UK cohort, we assess the consistency of non-cognitive skills over a 17-year period throughout childhood and adolescence, their genomic architecture, and their associations with socioeconomic outcomes. We find that longitudinal measurement consistency is high for behavioural and communication skills but low for other non-cognitive skills, implicating a high noise to signal ratio for many non-cognitive skills. Consistent non-zero heritability estimates and genetic correlations applied to cross-sectional measures are observed only for behavioural difficulties. When aggregating across multiple measurements, we find evidence of low heritability 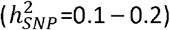 for behaviour, communication, self-esteem and locus of control. We find weak correlations between aggregate measures of skills, further supporting cross-sectional measurement error in the non-cognitive measures. Associations between non-cognitive skills and educational outcomes are observed for skills measured in mid to late childhood and these are at most a third of the size of IQ-education associations. These results suggest that individual measures designed to capture non-cognitive skills may be subject to considerable measurement error and provide unreliable indicators of children’s skills. However, aggregate measures that leverage longitudinal data may more reliably identify underlying non-cognitive traits.

## Introduction

Non-cognitive (or socioemotional) skills have been posited as an important driver of individual differences for a range of health and socioeconomic outcomes ^1–8^. Non-cognitive skills are broadly defined as “personality traits, goals, character, motivations, and preferences” ^3^ an are considered complementary to cognitive measures such as intelligence or general cognitive ability ^8^. A wide range of characteristics have been proposed to fall under the umbrella term of non-cognitive skills, including multifaceted personality and behavioural traits such as persistence, motivation and temperament ^2,8^.

Research into noncognitive skills has demonstrated the role that these may play in outcomes. Borghans and colleagues reported that personality explains 16% of the variance in achievement scores in a US sample ^7^. A meta-analysis found modest correlations between conscientiousness and academic achievement (*r*=0.24) but not for other personality types (r=−0.05 to 0.6) ^9^. Schmidt and Hunter showed that integrity and conscientiousness predict job and training performance ^10^. Von Stumm and colleagues show that intellectual curiosity and effort are associated with academic attainment ^11^.

The evidence supporting the role of non-cognitive skills on socioeconomic and health outcomes is however diverse and inconsistent. While research has demonstrated that associations between personality types and life outcomes replicate well ^12^, a recent systematic review showed that many other non-cognitive skills have small and heterogenous effects ^13^. These heterogenous effects may be driven by poor measurement of non-cognitive skills. Many studies have been restricted by short follow-up ^13^ or a lack of longitudinal data on non-cognitive skills to measure measurement consistency over time ^14^. Test-retest reliability has been estimated at 0.31 for personality in childhood ^15^, 0.12 and 0.46 to 0.61 for risk aversion, and 0.49 for locus of control ^16,17^. These values are lower than those reported for cognitive skills (c.f. 0.52 for digit span to 0.82 for reading) ^16^, suggesting that non-cognitive skills may be more variable or have higher measurement error than cognitive skills ^3^.

Few studies have measured a wide battery of non-cognitive skills, but modest correlations have been observed between some non-cognitive skills including grit and conscientiousness ^18^; academic effort and academic problems ^6^; and internalising and externalising behaviours ^19^. While modest ^6,18^, these correlations support a broad multidimensional definition of non-cognitive skills. Due to data limitations, many studies have also been unable to adjust estimates for cognitive ability despite its strong attenuating effect on non-cognitive associations with outcomes ^10,11^. More evidence is required about the relationship between non-cognitive skills in childhood and later outcomes, and the factors that mediate these effects.

The measurement consistency of non-cognitive skills and the robustness of their associations with socioeconomic outcomes can also be explored with genetic data. Identifying signal in genetic analyses of a phenotype requires both a genotypic component and reliable measurement of the phenotype. Personality types have been the most studied of the non-cognitive skills with heritability estimated at ^~^0.5 ^20^. Heritability estimates for other non-cognitive skills has varied widely from 0.44 to 0.79 for self-control ^21,22^; 0.36 for alienation ^6^; 0.83 for academic effort ^6^; 0.31 to 0.56 for aspects of openness ^23^; 0.18 to 0.49 for aspects of conscientiousness ^24^; and 0.40 for enjoyment and self-perceived ability ^25^. Sibling correlations for non-cognitive skills have also been observed ^26^. There is also genetic evidence that personality types are stable over time ^27^, providing support that personality measures are sufficiently free from measurement error to isolate genetic signal. There has however not yet been a comprehensive analysis investigating the genetics of multiple non-cognitive skills within the same sample of individuals ^28^.

In this study, we contribute to the literature with a comprehensive analysis into the phenotypic and genotypic measurement consistency of non-cognitive skills. Using data from a UK cohort study, we investigate the relationships between 10 non-cognitive skills, educational achievement and labour market outcomes. We answer two related research questions: 1) How consistent are non-cognitive skills measured in a large cohort study over time? 2) Do non-cognitive skills associate with socioeconomic outcomes once other factors including cognitive ability have been accounted for?

## Materials and Methods

### Study sample

Participants were children from the Avon Longitudinal Study of Parents and Children (ALSPAC). Pregnant women resident in Avon, UK with expected dates of delivery 1st April 1991 to 31^st^ December 1992 were invited to take part in the study. Two phases of recruitment resulted in a total sample of 14,899 children who were alive at one year of age, of whom 7,988 had genetic data. For full details of the cohort profile and study design see ^29,30^. The ALSPAC cohort is largely representative of the UK population when compared with 1991 Census data; there is under representation of some ethnic minorities, single parent families, and those living in rented accommodation ^29^. Ethical approval for the study was obtained from the ALSPAC Ethics and Law Committee and the Local Research Ethics Committees. We used the largest available samples in each of our analyses to increase precision of estimates, regardless of whether a child has data on other non-cognitive skills (see Supplementary Table 1 for sample sizes).

### Genetic data

DNA of the ALSPAC children was extracted from blood, cell line and mouthwash samples, then genotyped using references panels and subjected to standard quality control approaches. Briefly, ALSPAC children were genotyped using Illumina HumanHap550 quad chip genotyping platforms and ALSPAC mothers were genotyped using Illumina human660W-quad array platforms. All individuals with non-European ancestry were removed and exclusions were made based on gender mismatches; minimal or excessive heterozygosity; disproportionate levels of individual missingness (>3%); insufficient sample replication (IBD < 0.8); low SNP frequency (<1%), call rate (<95%) and violations of Hardy-Weinberg equilibrium (P < 5E-7). Genotypes in common were combined and imputed to the Haplotype Reference Consortium (HRCr1.1, 2016) panel of approximately 31 000 phased whole genomes, giving 8,237 eligible children with available genotype data after exclusion of related individuals using cryptic relatedness measures. Principal components were generated by extracting unrelated individuals (IBS < 0.05) and independent SNPs with long range LD regions removed. For full details of genotyping see the Supplementary Material.

### Non-cognitive skills

We used all non-cognitive skills that were available in ALSPAC except for attention which was omitted due to low responses. The supplementary material contains more detailed information on the measures used, their components and a timeline of when all skills and outcomes were measured (Supplementary Figure 1).

### SDQ

Study mothers reported on the Strengths and Difficulties Questionnaire (SDQ) for children on seven occasions at child ages 4, 7, 8, 10, 12, 13 and 16 years, and the children’s teachers completed an SDQ questionnaire for each child at ages 7 and 10. The SDQ scale is used to assess child emotional and behavioural difficulties, consisting of 25 items that cover common areas of emotional and behavioural difficulties. Total SDQ score was defined as the count of problems on the first four scales. Sensitivity analyses were run using the internalising and externalising sub-scales. All SDQ scores are reverse coded so that high values refer to fewer problems.

### Denver

The Denver Developmental Screening Test was used to identify developmental problems in young children at ages 6, 18, 30 and 42 months. ALSPAC mothers reported on their child’s development in response to 42 questions across four different categories: social and communication skills, fine motor skills, hearing and speech, and gross motor skills. Prorated scores combining all four scales were used with missing values assigned the mean score of that child’s responses, provided that three or less items had missing scores.

### Social skills

Social skills at age 13 were determined using a battery of 10 questions reported by the study mother. Responses were reported on a five-point scale and then summed to provide a total overall social skills score.

### Communication

Communication at 6 months was calculated from mother-reported responses to a battery of eight questions asking about the development of their child’s communication skills. At age 1 communication was calculated from mother-reported responses using response to 82 questions on the MacArthur Infant Communication questionnaire. At 18 months communication was calculated using mother-reported responses to a battery of 14 questions asking about the development of their child’s communication skills. At age 3 communication was calculated from mother-reported responses to a battery of 123 questions forming a vocabulary score. At age 10 communication was calculated from mother-reported responses as the sum of five domains of communication from a total battery of 39 questions.

### Self-esteem

Self-esteem at age eight was measured using self-report responses to the 12-item shortened form of Harter’s Self Perception Profile for Children comprising the global self-worth and scholastic competence subscales. Self-esteem at age 18 was measured using self-report responses to 10 questions of the Bachman revision of the Rosenberg Self-Esteem Scale.

### Persistence

Persistence at age 6 months was measured as a weighted score from mother-reported responses to seven questions relating to child temperament. At age 2 persistence was measured as a weighted score from nine mother-reported responses. At age 7 persistence was recorded as the participants persistence when completing tasks under direct assessment.

### Locus of control

Locus of control, the strength of connection between actions and consequences, was measured at age eight using responses to 12 questions from the shortened version of the Nowicki-Strickland Internal-External (NSIE) scale. At age 16 it was measured using the 12 item Nowicki-Strickland Locus of Control Scale.

### Empathy

Empathy was measured at age seven using mother reported responses to five questions about the child’s attitudes towards sharing and caring.

### Impulsivity

Impulsivity was measured during two sessions at the age 8 direct assessment using a behaviour checklist. Testers rated whether the children demonstrated restlessness, impulsivity, fleeting attention, and lacking persistence. At age 11 the children were asked a battery of 10 questions designed to capture impulsive behaviour.

### Personality

Personality was measured at age 13 using the five-factor model of personality to capture five broad and independent dimensions of personality; the “Big Five” (extraversion, neuroticism, agreeableness, conscientiousness, and intellect)^31^. These were measured using self-report responses to 50 items of the International Personality Item Pool.

### Cognitive skills and outcomes

#### IQ

Intelligence was measured during the direct assessments at ages eight and 15 using the short form Wechsler Intelligence Scale for Children (WISC) from verbal, performance, and digit span tests and the Wechsler Abbreviated Scale of Intelligence (WASI) from vocabulary and matrix reasoning tests respectively. These assessments were administered by members of the ALSPAC psychology team overseen by an expert in psychometric testing. The short form tests have high reliability and the ALSPAC measures utilise subtests with reliability ranging from 0.70 to 0.96.

#### Educational achievement

We used four measures of educational achievement. The first three were average fine-graded point scores from three end of ‘Key Stage’ assessments during compulsory education at ages 11, 14 and 16. The fourth measure was a ranking of grades attained in post-compulsory A-levels at age 18, which are required for progression to university education. We used a measure of the three highest A-level grades grouped into ordered categories (see ^32^ for a detailed description). At the time the cohort were studying, A-levels were non-compulsory and therefore all participants who did not continue into further education were set to missing. All measures were obtained through data linkage to the UK National Pupil Database (NPD) which represents the most accurate record of educational achievement available in the UK. All education data were extracted from the NPD Key Stage 4 (age 16) and Key Stage 5 databases (for further information see https://www.gov.uk/government/collections/national-pupil-database).

#### Employment

At age 23 participants were asked to report whether they were in full time paid employment of more than 30 hours per week, with responses coded as binary.

#### Not in education employment or training (NEET)

Because some participants may not be employed because they are still in full-time education or training, we used a measure of not in education employment or training (NEET) at age 23 with responses coded in binary. The use of a NEET measure ensures that employment results are not biased by participation of the cohort in education or training.

#### Income

At age 23 participants were asked to report their take-home pay each month if they reported being in paid employment, with responses banded into the following categories: £1-£499; £500-£999;

£1000-£1499; £1500-£1999; £2000-£2499; £2500-£3000; £3000+.

#### Non-response

We also include a binary measure of questionnaire non-response at ages 18 and 24 to allow us to investigate correlations between non-cognitive skills and cohort participation.

### Statistical analysis

We estimated phenotypic correlations between each measurement-pair (45 measurements [34 non-cognitive skills; 2 cognitive skills; 7 socioeconomic outcomes; 2 non-response measures]; 990 unique measurement-pairs). Heritability of each occasion-specific non-cognitive skill was estimated using genomic-relatedness-based restricted maximum likelihood (GREML) in the software package GCTA (see ^33^ for a detailed description of this method). GCTA uses measured SNP level variation across all SNPs to estimate the genetic similarity between each pair of unrelated individuals in the sample. Univariate analyses are specified as:

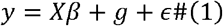

where *y* is the inverse normally rank transformed sex and age of measurement standardised measure of phenotype, *X* is a series of covariates indicating the first 20 principal components of inferred population structure to control for systematic differences in allele frequencies due to ancestral differences between different subpopulations (population stratification), *g* is a normally distributed random effect with variance 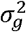 denoting the contribution of SNPs, and *ϵ* is residual error with variance 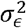. Heritability is then defined as the proportion of phenotypic variance that can be statistically explained by common genetic variation while holding inferred population structure constant:

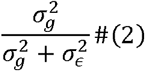

To estimate the extent to which non-cognitive traits share underlying genetic architecture we estimate genetic correlations between each phenotype-pair. Genetic correlations provide an estimate of the proportion of variance that two phenotypes share due to genetic variation (the overlap of genetic associations between two phenotypes). Genetic correlations are estimated as:

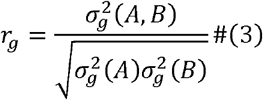

Where *r*_*g*_ is the genetic correlation between phenotypes *A* and *B*, 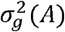 is the genetic variance of phenotype *A* and 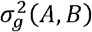 is the genetic covariance between phenotypes *A* and *B*. All analyses are adjusted for false discovery rate using the Benjamini-Hochberg procedure ^34^ and include the 20 principal components of population structure.

## Results

### How consistent are non-cognitive skills over time?

Behavioural skills measured on the SDQ scale were consistent over time (parent reported SDQ *r*=0.36 to 0.73; teacher reported SDQ *r*=0.28 to 0.53), though consistency decreased with greater elapsed time between measures increased (Figure 1). Correlations between parent reported and teacher reported SDQ measures were weak even when measured contemporaneously. Correlations for other non-cognitive measures over time were generally low (Figure 1), suggesting within-trait inconsistency (highly variable skills) or lack of reliability (high measurement error). Correlations amongst the Big 5 personality types were low except between the intellect/imagination and agreeableness subscales (*r*=0.46). While the phenotypic correlations for repeat measures of many non-cognitive skills were low, they were generally positive. The weak temporal phenotypic correlations of non-cognitive measures contrast sharply to those for cognitive measures of IQ (*r*=0.60) and measures of educational achievement (*r*>0.78 for compulsory education).

**Figure 1:**
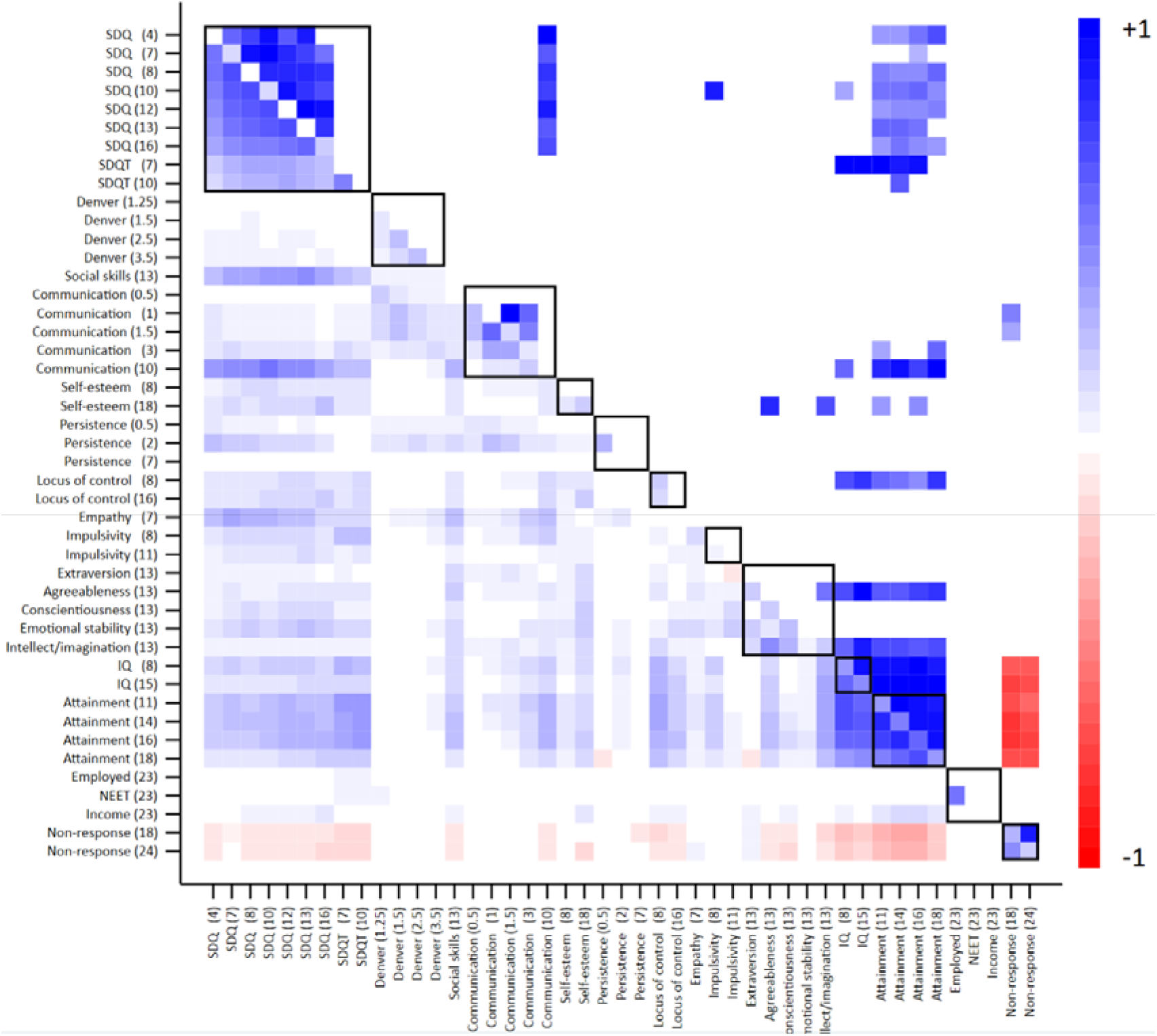
Heritability, phenotypic correlations and genetic correlations of skills and outcomes. Age in parentheses. SDQ, strengths and difficulties; NEET, not in education, employed or training. Values on the diagonal represent univariate heritabilities; values below the diagonals represent phenotypic correlations; values above the diagonal represent genetic correlations. Multiple testing was handled using an FDR threshold of 5%. Empty cells display correlations that fell below FDR threshold. Black outline boxes indicate the same skills measured at different occasions. See Supplementary Tables 2 & 3 for full estimates.

### How correlated are different non-cognitive skills?

Phenotypic correlations between measures of different non-cognitive skills were generally low (Figure 1). Parent-reported and teacher-reported SDQ was the only measure that correlated consistently with other non-cognitive skills at *r*>0.2. This between-trait correlation was strongest for social skills (*r* =0.24 to 0.49), communication at age 10 (*r* =0.29 to 0.56) and empathy (*r* =0.17 to 0.38). Between-skill phenotypic correlations were almost exclusively positive, suggesting that where patterns were observed, children who scored high on one non-cognitive skill generally also scored highly on another. Teacher reported measures of SDQ, locus of control at age 8 and the intellect/imagination personality type correlated modestly with cognitive skills (*r*>0.3 for IQ at age 8). To further explore if measurement imprecision may be driving these low correlations, we took the mean value of non-cognitive skills that had been measured more than once (Figure 2). Weak correlations were observed between SDQ, Teacher SDQ, communication, self-esteem, locus of control and impulsivity. Positive correlations between these mean variables and educational achievement were observe for all variables other than persistence.

**Figure 2:**
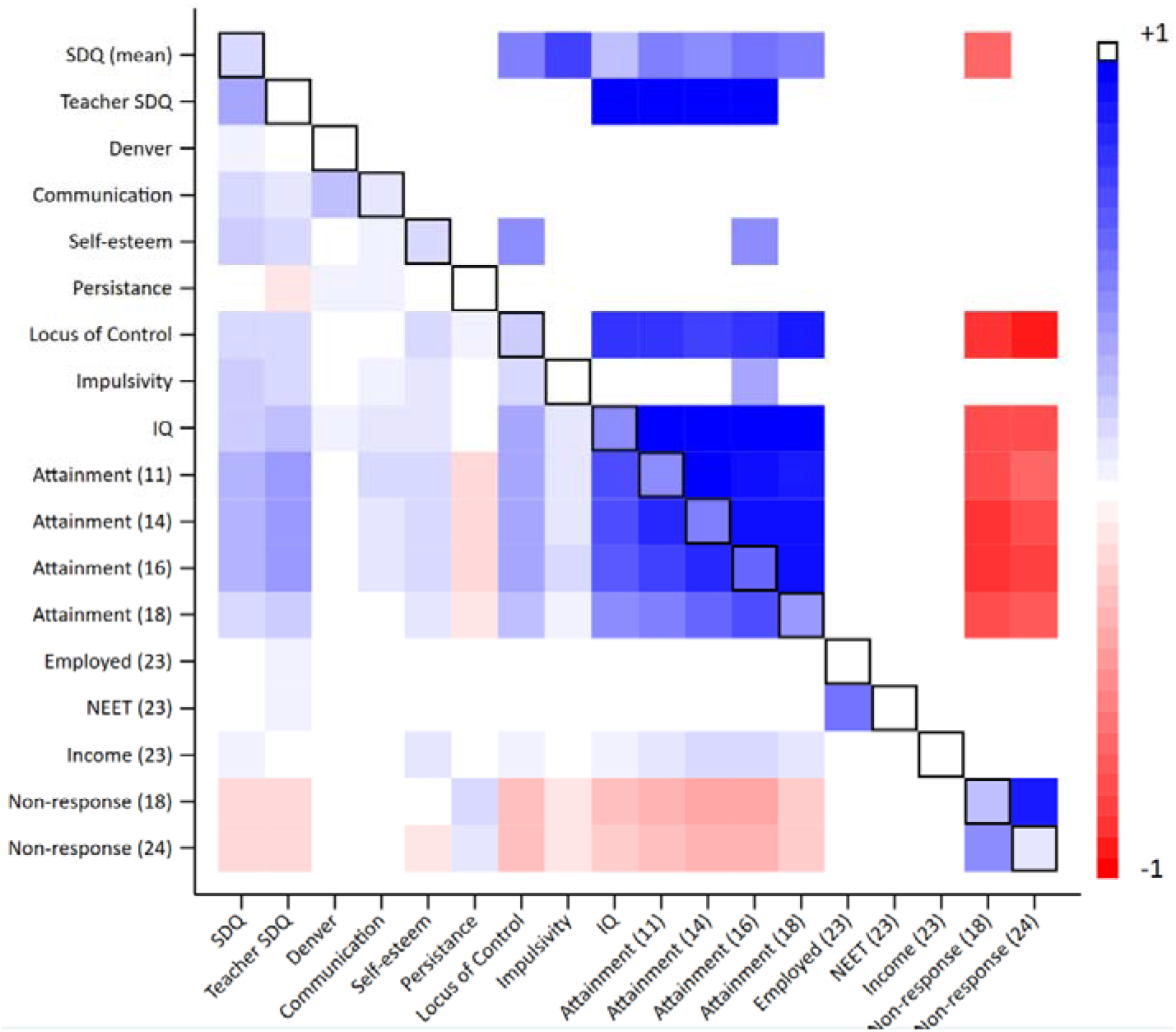
Heritability, phenotypic correlations and genetic correlations of aggregate skills and outcomes. Age in parentheses. SDQ, strengths and difficulties; NEET, not in education, employed or training. Values on the diagonal represent univariate heritabilities; values below the diagonals represent phenotypic correlations; values above the diagonal represent genetic correlations. Multiple testing was handled using an FDR threshold of 5%. Empty cells display correlations that fell below FDR threshold. Black outline boxes indicate the same skills measured at different occasions. See Supplementary Tables 4 & 5 for full estimates.

### Do non-cognitive skills associate with education and labour market outcomes?

Phenotypic correlations with educational achievement were highest for the SDQ scale (*r*=0.13 to 0.42) (Figure 1). The teachers’ SDQ assessments were more strongly associated than the contemporaneous parent reports. For example, the correlation between educational achievement at age 11 and teacher reported SDQ at age 10 was *r*=0.40 while the correlation with parent reported SDQ at age 10 was *r*=0.29. Correlations with educational outcomes were modest for social skills (*r*=0.13 to 0.27); communication at age 10 (*r*=0.18 to 0.34), locus of control (*r*=0.26 to 0.36 at age 8; *r*=0.19 to 0.28 at age 16), the agreeableness and the intellect/imagination subscales of the Big 5 (*r*=0.16 to 0.28 and *r*=0.27 to 0.39 respectively). Correlations between educational achievement and cognitive ability were considerably higher (*r*=0.43 to 0.73). Phenotypic correlations for non-cognitive skills with employment, NEET and income were low (*r*<=0.08 for employment; *r*<=0.07 for NEET; *r*<=0.19 for income). Correlations between labour market outcomes and cognitive skills were also very low (*r*<=0.09). Non-cognitive and cognitive skills were generally weakly negatively correlated with non-response at ages 18 and 24 (*r*=0.05 to 0.15). Only the extraversion subscale of the Big 5 was positively correlated with non-response (*r*=0.06).

To further investigate the potential impact of skills we ran a series of multivariable regressions of age 16 educational achievement on non-cognitive and cognitive skills (Figure 3). Each skill was standardised and analysed independently controlling for sex, month of birth and IQ at age 8. A one standard deviation (SD) increase in non-cognitive skill measures was associated with a 0.04 SD decrease to 0.21 SD increase in age 16 achievement. There was considerable heterogeneity for estimates between skills and measurement occasions for many non-cognitive measures. Associations were generally larger the closer in time non-cognitive skills were measured to educational achievement. By comparison, a one SD increase in cognitive skills was consistently associated with a 0.5 SD increase in achievement. These patterns were similar for educational achievement at all ages (Supplementary Figure 2).

**Figure 3:**
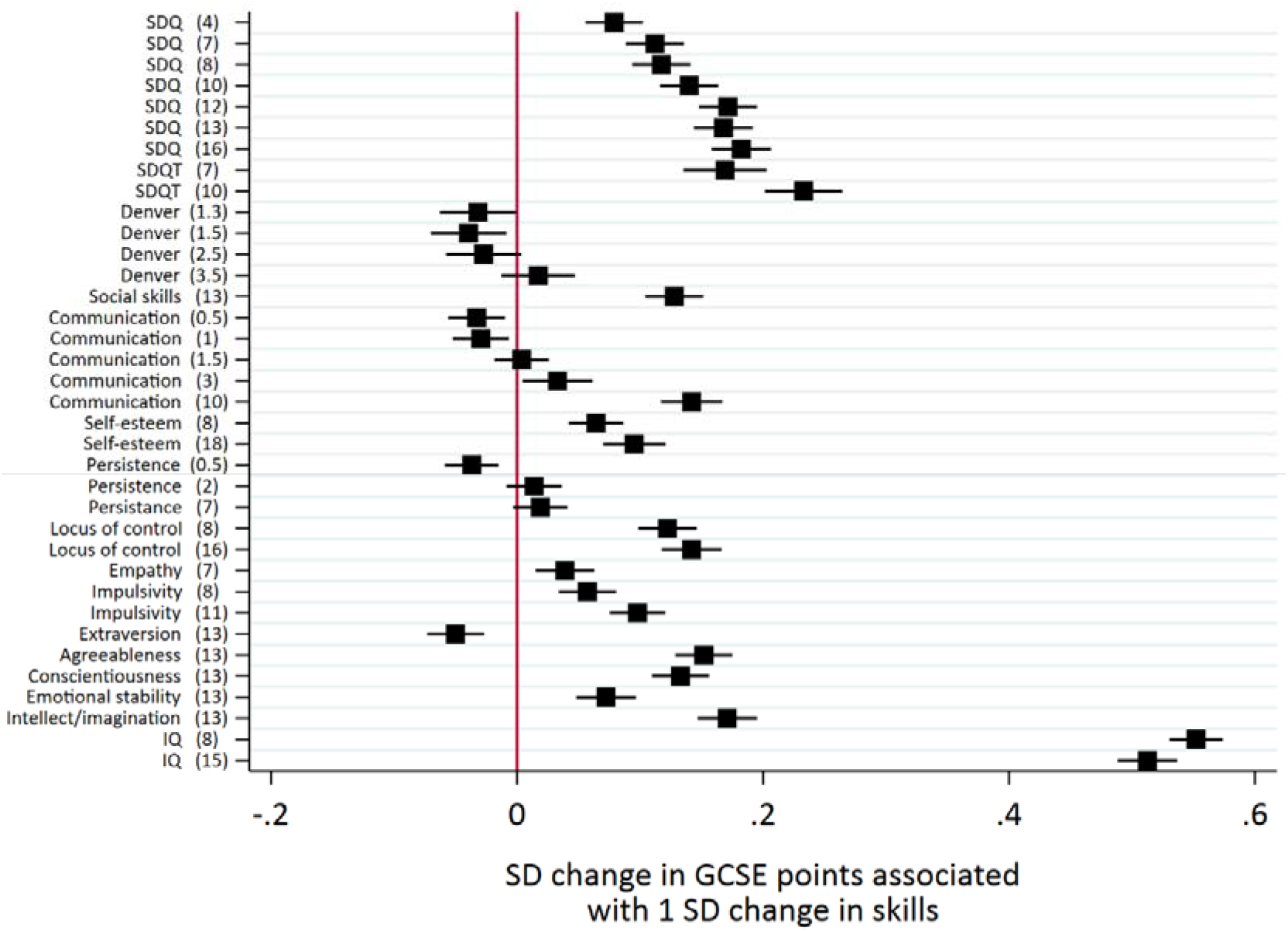
Associations between skills and educational attainment adjusted for sex, month of birth and IQ at age 8. Educational attainment measured as exam point score at age 16. Age (years) in parentheses. SDQ, strengths and difficulties.

### Are non-cognitive skills heritable?

Figure 4 displays the heritability estimates for all our measures. The parent reported SDQ scale was the only non-cognitive measure for which there was reasonable evidence of non-zero heritability at multiple occasions (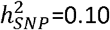 to 0.23). Results were broadly comparable when using the internalising and externalising subscales of the SDQ (Supplementary Figure 3). Non-zero heritability was also observed for communication at age 18 months 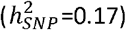, self-esteem at age 18 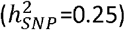, locus of control at age 8 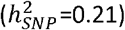, and the intellect/imagination subscale of personality type 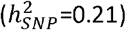. There was imprecision in the heritability point estimates, but the upper limit of the 95% confidence intervals were estimated below 0.3 for most non-cognitive skills. Heritability of the mean value non-cognitive skills was estimated with greater precision, but there was evidence of heritability for the mean responses of SDQ, 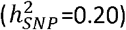, communication 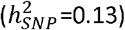, self-esteem and locus of control 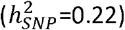 (Figure 5). The heritability of cognitive skills was higher at 0.43 at age 8 and 0.47 at age 15. Educational outcomes were highly heritable 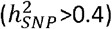 but there was little evidence of heritability for the labour market outcomes. Heritability of questionnaire non-response was estimated higher and with less uncertainty than the non-cognitive measures (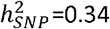 and 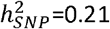 at ages 18 and 24 respectively).

**Figure 4:**
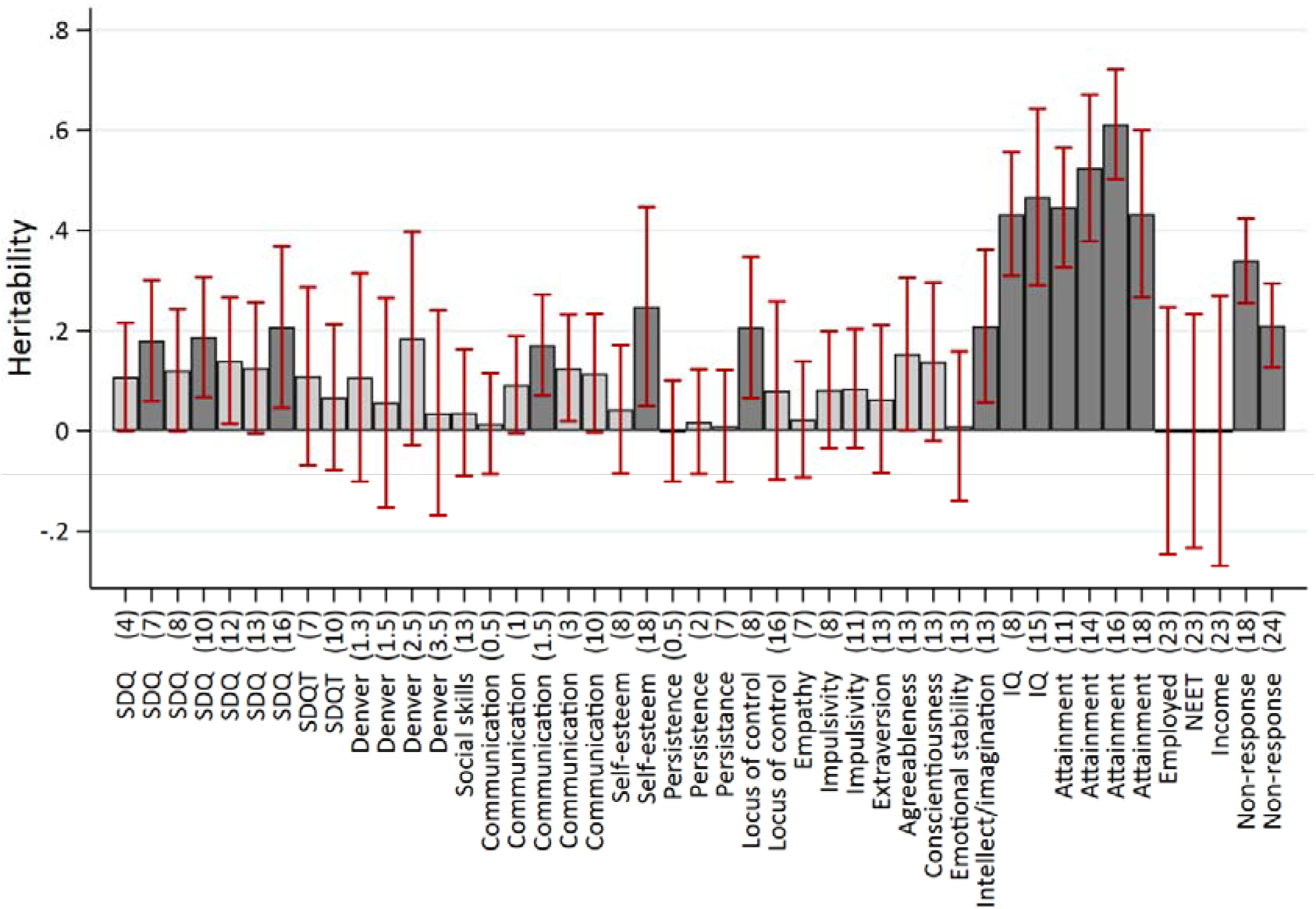
Heritability of skills and outcomes. Age in parentheses. SDQ, strengths and difficulties; NEET, not in education, employed or training. Dark shaded bars represent estimates below FDR threshold. Heritability estimated using GCTA-GREML on the full sample available for each phenotype. See Supplementary Table 1 for full estimates.

**Figure 5:**
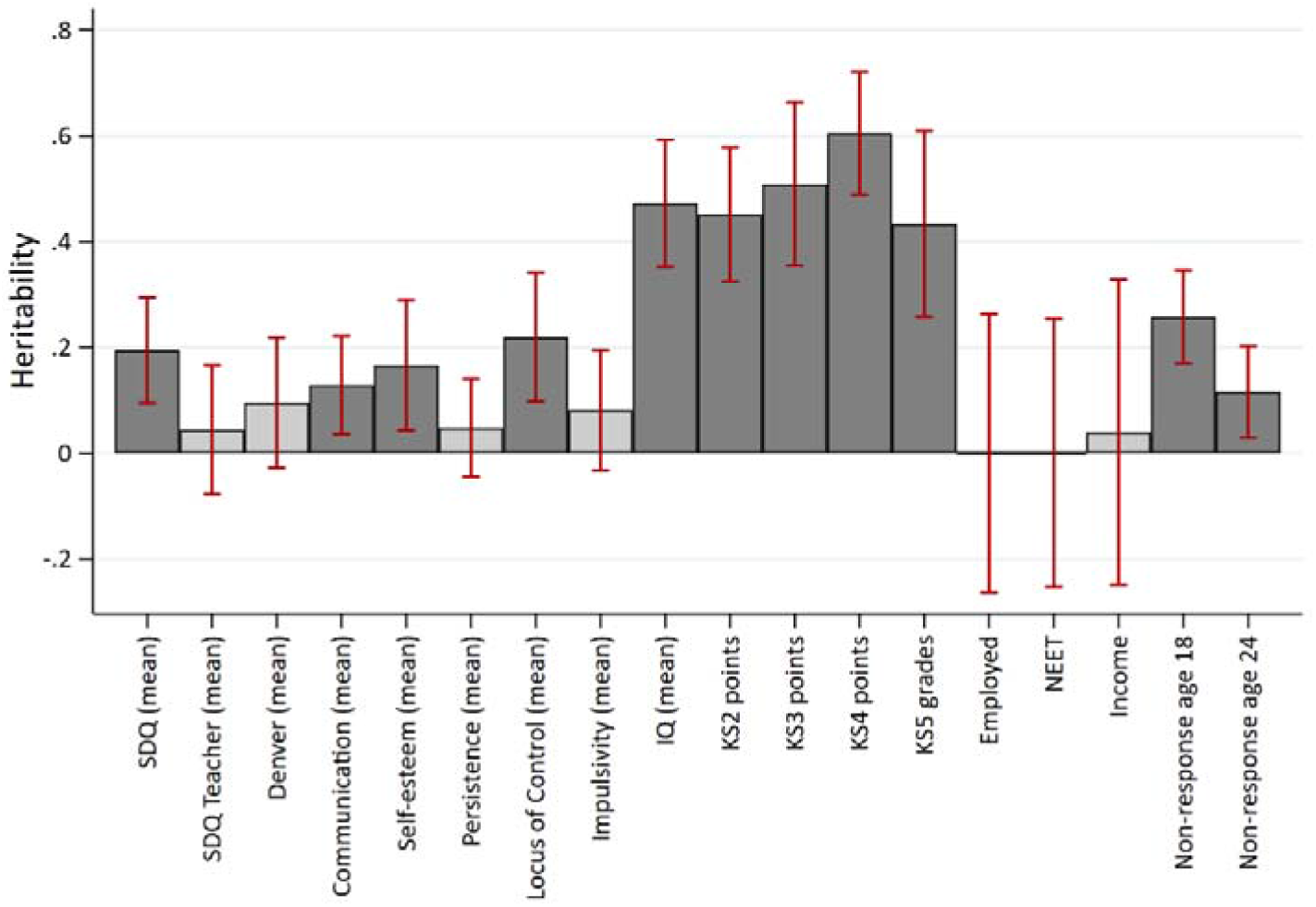
Heritability of aggregate skills and outcomes. Age in parentheses. SDQ, strengths and difficulties; NEET, not in education, employed or training. Dark shaded bars represent estimates below FDR threshold. Heritability estimated using GCTA-GREML on the full sample available for each phenotype. See Supplementary Table 1 for full estimates.

### Do non-cognitive skills have a shared genetic architecture?

There was strong evidence for within trait genetic correlations over time for only the parent reported SDQ measures (*rG*= 0.62 to 1.00) and IQ, which had near-unity genetic overlap (*rG*= 0.97). There was very limited evidence for genetic correlations across different non-cognitive skills (Figure 1, above the diagonal), though estimation precision was low for most skills given the low SNP heritabilities. Genetic correlations between different non-cognitive measures were only observed between the parent reported SDQ measures and communication at age 10 (*rG*=0.68 to 0.91). This remained the case when using mean values of the non-cognitive skills (Figure 3); genetic correlations were only observed between SDQ and locus of control (*r*=0.50), SDQ and impulsivity (*r*=0.76), and self-esteem and locus of control (*r*=0.50). The SDQ measures, communication at age 10, self-esteem at age 18, the agreeableness and the intellect/imagination subscales of the Big 5 personality types had strong genetic correlations with educational achievement. These were higher for teacher report at age 7 than any of the parents and teacher reported at age 16 SDQ measures. There was strong evidence for non-zero genetic correlations with IQ and education for the mean values of parent SDQ, teacher SDQ and locus of control. Genetic correlations between cognitive measures and educational achievement are all estimated near unity (*rG*>0.96). There was little evidence of genetic correlations between labour market outcomes and any of the non-cognitive or cognitive skills.

## Discussion

Our results provide longitudinal evidence into the consistency of a large range of non-cognitive skill measures throughout childhood and adolescence. Measures of behavioural and communication skills were correlated phenotypically and genotypically over time. Correlations of other measured non-cognitive skills over time were low, contrasting with previous research that has evidenced temporal stability of non-cognitive skills ^14,15^. This heterogeneity may be due to the diversity of non-cognitive skills measured. For example, the SDQ scale used responses to a large battery of validated questions which are likely to be more reliable than other non-cognitive skill measures used here. The increased estimation precision that we observed when using aggregated values of non-cognitive skills supports this. By incorporating information from multiple occasions, aggregate measures are expected to contain less measurement error than individual measures. Future research proposing more detailed measurement of non-cognitive skills may therefore improve the ability of measures to capture underlying skills. The lack of consistency may have also reflected genuine temporal intra-individual variation in the expression of skills or differences over time (e.g. due to schooling). It is important to note there is some variation in the measurement of skills that we used. This is somewhat unavoidable as skill measurements vary depending on the age at which a child is assessed, but measurement artefact may nevertheless reflect some of the variation in measurement. Non-cognitive skills have been argued as promising targets for interventions to improve outcomes because they are potentially more modifiable than cognitive skills ^1,3,15^. However, ideally to be suitable for policy interventions, non-cognitive measures must reliably pick up a consistent signal of an underlying skill. While changes to non-cognitive skills are required (and indeed anticipated) during interventions, they should remain relatively stable in individuals that experience no intervention. These findings suggest that many measures of non-cognitive skills may be too inaccurate for accurately measuring the impact of interventions.

Correlations between different non-cognitive skills were weak, supporting the notion that these constructs capture empirically different psychosocial phenomena rather than a single underlying or latent non-cognitive factor. The SDQ scale was the only measure to consistently correlate with other non-cognitive skills, conforming to a previous study that found low phenotypic correlations between different non-cognitive skills ^6^. Stronger between-trait correlations were observed using aggregated values of the non-cognitive skills, which again may reflect reduced measurement error. The cognitive measures we used were more strongly correlated over the same period, though this may reflect that they are more reliably measured than the non-cognitive measures. Future studies that combine longitudinal data on non-cognitive skills with multi-source multi-method approaches may benefit from greater measurement precision.

We found limited evidence of genetic correlations within or between non-cognitive skills, with only the parent reported SDQ demonstrating consistent genetic architecture over time. This may reflect the influence of shared parent-offspring genetics on reporting, or parental genetics influencing offspring non-cognitive skills indirectly through dynastic effects ^35^ (there would be no such shared teacher-student genetics). Previous twin studies have found strong genetic correlations between non-cognitive and cognitive skills ^23,24^, but our results did not support this. The differences between these results and previous studies may have arisen due to differences in the quality of non-cognitive skill measurements; differences due to cross-sectional and longitudinal study designs; reporting bias by the study mothers (for example where their child performed poorly); differences in the study populations; selective reporting and the use of samples or measures of convenience in previous studies ^13^. While we reported a range of non-cognitive skills in ALSPAC, many previous studies have reported only one or a small number of non-cognitive skills.

Our results also showed that associations between non-cognitive skills and socioeconomic outcomes were generally weak, contradicting findings from previous individual studies ^1–3^ but supporting a recent systematic review ^13^. Behavioural problems as captured by the SDQ scale, social skills, and locus of control were the only non-cognitive measures to phenotypically associate with educational outcomes strongly and consistently. Associations were stronger between educational achievement and the teacher than the parent reported measures of the SDQ scale. This may suggest that teachers more objectively identify education related problematic behaviour than parents, or that teachers’ response to their pupils directly influences their educational outcomes. Non-cognitive skills capture a diverse range of characteristics and may have varying relevance to educational and labour market outcomes; further longitudinal research is required to better elucidate these relationships. Neither non-cognitive nor cognitive skills associated strongly with the labour market outcomes measured here. While this contrasts with previous studies ^1,3^, our labour market outcomes were observed soon after entry to the labour market and therefore may have more closely resembled institutional effects than non-cognitive skills. Some have argued that non-cognitive skills and personality traits are as important as cognitive skills for many dimensions of behaviour and socioeconomic outcomes ^5^, but our results do not support this. The standardized effect sizes of non-cognitive skills were at most a third that of cognitive skills.

Many non-cognitive and cognitive skills were weakly negatively correlated with non-response phenotypically, suggesting that individuals who scored low on these skills were more likely to later drop out of the ALSPAC study. This may have important implications for participant representativeness and generalisability in cohort studies. Cognitive ability and achievement were negatively genotypically correlated with non-response, adding to the growing body of evidence that non-response is genetically patterned ^36^.

The SDQ scale was the only non-cognitive measure for which we consistently found strong evidence of heritability (^~^0.15), far lower than the estimated heritabilties for cognitive measures (^~^0.45) and educational achievement (^~^0.50). Heritability and stability depend on the developmental period being examined and so comparison of estimates is difficult. Previous research has demonstrated that the heritability of cognitive ability rises with age while the heritability of personality traits decreases with age ^37^, which our data contradicts. Many of the genetic correlations we observed were imprecise due to the low heritability estimates, but our estimates suggested an upper bound heritability of 0.3 for most non-cognitive skills.

This study has several limitations. First, it is possible that measurement error was unusually high in the non-cognitive measures used in the ALSPAC study. However, the measures used in ALSPAC have been widely validated are consistent with those used in previous studies ^19^. Furthermore, measurement error would need to have been high across all measures used from birth to age 18 so to explain these results. Our findings were consistent when using aggregated measures of non-cognitive skills where they had been measured at least twice, suggesting that our results were not driven by differential measurement error across occasions. Future studies into test-retest reliability of non-cognitive skills based on different longitudinal samples could help resolve these questions. Second, many of the genetic correlations were estimated with extremely low precision, often being constrained at the values of −1 or 1 (see supplementary Table 2). This is likely due to the low estimated heritability of the non-cognitive skills. Low heritability implies a small genotypic contribution (either in the number of individual variants associated or the strength of associations) and therefore lower power to detect genetic correlations between these skills. Our sample sizes are fairly low to detect such small univariate and bivariate genetic associations. Future studies conducted on larger samples are required to more accurately the estimate heritability of, and genetic correlations between non-cognitive skills and other phenotypes. Third, our genetic associations may have been biased by uneven linkage disequilibrium, residual population structure, or assortative mating ^35,38^. We controlled for the first twenty principle components of population structure to account for population structure but this may not have accounted for all differences ^39^. Assortative mating is thought to be low for non-cognitive traits ^20^, but will have inflated our genetic associations if present ^40^. It is possible that assortment on non-cognitive skills may be negative and future work is required to determine this. Fourth, the use of labour market outcomes at age 23 may mean that some of our study participants have not yet transitioned into their stable career employment. While our use of a NEET outcome variable reduces the problem that those still in education or unemployed have not yet entered the labour market, it does not provide any indication that employed study participants have entered desired or long-term employment. Finally, it is possible that some of the measures could reflect parental rather than child genes where they are parent rather than self-reported.

In conclusion, our results highlight that non-cognitive skills are likely to be highly heterogenous. Measures of noncognitive skills were varied both over time and across different measures in the same individuals, but some individual measures such as the SDQ demonstrated strong internal consistency throughout childhood. Furthermore, many non-cognitive measures associated weakly with educational and employment outcomes at entry to the labour market, particularly when measured early in childhood.

## Acknowledgements

We are extremely grateful to all the families who took part in this study, the midwives for their help in recruiting them, and the whole ALSPAC team, which includes interviewers, computer and laboratory technicians, clerical workers, research scientists, volunteers, managers, receptionists and nurses. The Medical Research Council (MRC) and the University of Bristol support the MRC Integrative Epidemiology Unit [MC_UU_12013/1, MC_UU_12013/9, MC_UU_00011/1]. A comprehensive list of grants funding is available on the ALSPAC website (http://www.bristol.ac.uk/alspac/external/documents/grant-acknowledgements.pdf); data used at age 23 was specifically funded by the Wellcome Trust and MRC [102215/2/13/2]. Study data were collected and managed using REDCap electronic data capture tools hosted at the University of Bristol. REDCap (Research Electronic Data Capture) is a secure, web-based application designed to support data capture for research studies, providing 1) an intuitive interface for validated data entry; 2) audit trails for tracking data manipulation and export procedures; 3) automated export procedures for seamless data downloads to common statistical packages; and 4) procedures for importing data from external sources. The study website contains details of all the data that is available through a fully searchable data dictionary and variable search tool at http://www.bristol.ac.uk/alspac/researchers/our-data/. The Economics and Social Research Council (ESRC) support NMD via a Future Research Leaders grant [ES/N000757/1] and TTM via a Postdoctoral Research Fellowship [ES/S011021/1]. The UK Medical Research Council and Wellcome (Grant ref: 102215/2/13/2) and the University of Bristol provide core support for ALSPAC. This publication is the work of the authors and Tim Morris will serve as guarantors for the contents of this paper. GWAS data was generated by Sample Logistics and Genotyping Facilities at Wellcome Sanger Institute and LabCorp (Laboratory Corporation of America) using support from 23andMe. No funding body has influenced data collection, analysis or its interpretations. This work was carried out using the computational facilities of the Advanced Computing Research Centre - http://www.bris.ac.uk/acrc/ and the Research Data Storage Facility of the University of Bristol - http://www.bris.ac.uk/acrc/storage/.

## Conflict of interest

The authors declare no conflict of interest.

## Notes

### Competing Interest Statement

The authors have declared no competing interest.

