## Supplementary Material for "Consistency of non-cognitive skills and their relation to educational outcomes in a UK cohort"

### Genetic data

DNA of the ALSPAC children was extracted from blood, cell line and mouthwash samples, then genotyped using references panels and subjected to standard quality control approaches. ALSPAC children were genotyped using the Illumina HumanHap550 quad chip genotyping platforms by 23andme subcontracting the Wellcome Trust Sanger Institute, Cambridge, UK and the Laboratory Corporation of America, Burlington, NC, US. The resulting raw genome-wide data were subjected to standard quality control methods. Individuals were excluded on the basis of gender mismatches; minimal or excessive heterozygosity; disproportionate levels of individual missingness (>3%) and insufficient sample replication (IBD < 0.8). Population stratification was assessed by multidimensional scaling analysis and compared with Hapmap II (release 22) European descent (CEU), Han Chinese, Japanese and Yoruba reference populations; all individuals with non-European ancestry were removed. SNPs with a minor allele frequency of < 1%, a call rate of < 95% or evidence for violations of Hardy-Weinberg equilibrium (P < 5E-7) were removed. Cryptic relatedness was measured as proportion of identity by descent (IBD > 0.1). Related individuals that passed all other quality control thresholds were retained during subsequent phasing and imputation. 9 115 individuals and 500 527 SNPs passed these quality control filters.

ALSPAC mothers were genotyped using the Illumina human660W-quad array at Centre National de Génotypage (CNG) and genotypes were called with Illumina GenomeStudio. PLINK (v1.07) was used to carry out quality control measures on an initial set of 10 015 individuals and 557 124 directly genotyped SNPs. SNPs were removed if they displayed more than 5% missingness or a Hardy-Weinberg equilibrium P value of less than 1.0e-06. Additionally, SNPs with a minor allele frequency of less than 1% were removed. Samples were excluded if they displayed more than 5% missingness, had indeterminate X chromosome heterozygosity or extreme autosomal heterozygosity. Samples showing evidence of population stratification were identified by multidimensional scaling of genome-wide identity by state pairwise distances using the four HapMap populations as a reference, and then excluded. Cryptic relatedness was assessed using a IBD estimate of more than 0.125 which is expected to correspond to roughly 12.5% alleles shared IBD or a relatedness at the first cousin level. Related individuals that passed all other quality control thresholds were retained during subsequent phasing and imputation. 9 048 individuals and 526 688 SNPs passed these quality control filters.

We combined 477 482 SNP genotypes in common between the sample of mothers and sample of children. We removed SNPs with genotype missingness above 1% due to poor quality (11 396 SNPs removed) and removed a further 321 individuals due to potential ID mismatches. This resulted in a dataset of 17 842 individuals containing 6 305 duos and 465 740 SNPs (112 were removed during liftover and 234 were out of HWE after combination). We estimated haplotypes using ShapeIT (v2.r644) which utilises relatedness during phasing. The phased haplotypes were then imputed to the Haplotype Reference Consortium (HRCr1.1, 2016) panel of approximately 31 000 phased whole genomes. The HRC panel was phased using ShapeIt v2, and the imputation was performed using the Michigan imputation server. This gave 8 237 eligible children and 8 196 eligible mothers with available genotype data after exclusion of related individuals using cryptic relatedness measures described previously. Principal components were generated by extracting unrelated individuals (IBS < 0.05) and independent SNPs with long range LD regions removed, and then calculating using the `--pca` command in plink1.90.

### Non-cognitive measures

#### SDQ

Study mothers reported on the Strengths and Difficulties Questionnaire (SDQ) for children on seven occasions at child ages 4, 7, 8, 10, 12, 13 and 16 years, and the children’s teachers completed an SDQ questionnaire for each child at ages 7 and 10. The SDQ is a scale used to assess child emotional and behavioural difficulties and is one of the most widely used questionnaires for evaluating psychological well-being amongst children. It consists of 25 items that cover common areas of emotional and behavioural difficulties (emotional symptoms; conduct problems; hyperactivity/inattention; peer relationship problems; and prosocial behaviour). Responses to each question are on a three-point scale: not true, somewhat true and certainly true, coded to scores of 0, 1 and 2 respectively. The individual subscales of the SDQ have relatively low prevalence in ALSPAC so we used total SDQ score which is defined as the count of problems on the first four scales. To ensure that our results are not being driven by differences in the internalising (emotional symptoms and peer relationship problems) or externalising (conduct problems and hyperactivity/inattention symptoms) subscales we also ran sensitivity analyses on these separate sub-scales. All SDQ scores are reverse coded so that high values refer to fewer problems.

#### Denver

The Denver Developmental Screening Test was used to identify developmental problems in young children at ages 6, 18, 30 and 42 months. ALSPAC mothers were asked to report their child’s development in response to 42 questions across four different categories: social and communication skills, fine motor skills, hearing and speech, and gross motor skills. Responses to questions were ‘often’, ‘once or twice’ and ‘not yet started’, and were coded with the values of 2, 1 and 0 respectively. Prorated scores combining all four scales were used to boost sample size with missing values assigned the mean score of that child’s responses, provided that three or less items had missing scores.

#### Social skills

Social skills at age 13 were determined using a battery of 10 questions reported by the study mother, such as being “easy to chat with, even if it isn’t on a topic that specially interests her”. Responses were reported on a five-point scale indicating the mother’s perception of how well her teenager’s social skills compared to her perception of the average teenager, and then summed to provide a total overall social skills score.

#### Communication

Communication at 6 months was calculated from mother-reported responses to a battery of eight questions asking about the development of their child’s communication skills such as “S/he turns towards someone when they are speaking”. At age 1 communication was calculated from mother-reported responses using the MacArthur Infant Communication questionnaire was derived at age one as the sum of mother reported responses to 82 questions across four domains of communication: understanding; vocabulary; non-verbal communication; and social development. At 18 months communication was calculated using mother-reported responses to a battery of 14 questions asking about the development of their child’s communication skills. At age 3 communication was calculated from mother-reported responses to a battery of 123 questions forming a vocabulary score. At age 10 communication was calculated from mother-reported responses as the sum of five domains of communication from a total battery of 39 questions. Responses were coded as 2, 1 and 0 reflecting the level of communication and understanding of communication that the child demonstrated. A final communication score was created as the sum of these responses.

#### Self-esteem

Self-esteem at age eight was measured using self-report responses to the 12-item shortened form of Harter’s Self Perception Profile for Children comprising the global self-worth and scholastic competence subscales. Self-esteem at age 18 was measured using self-report responses to 10 questions of the Bachman revision of the Rosenberg Self-Esteem Scale. At each age responses are summed to give an overall score. Responses are coded on a Likert scale with the value 0 assigned to the most negative statements and the value 4 assigned to the most positive statement, so that high scores correspond to high self-esteem.

#### Persistence

Persistence at age 6 months was measured as a weighted score from mother-reported responses to seven questions relating to child temperament. At age 2 persistence was measured as a weighted score from nine mother-reported responses. Questions included items such as “He perseveres for many minutes when working on a new skill (rolling over, picking up object, etc)” and “He loses interest in a new toy or game within an hour” and were recorded on a five-point Likert scale. At age 7 persistence was recorded by ALSPAC interview testers as the study child’s persistence when completing the word reading and decoding session. In this session children were shown a series of pictures accompanied by four words starting with the same letter as the picture and were asked to point to the correct word. Response options were coded into persistent (combining persistent and sometimes persistent) and non-persistent. Testers indicated whether they thought the child was persistent with tasks during the session. On all measures higher scores correspond to higher levels of persistence.

#### Locus of control

Locus of control, the strength of connection between actions and consequences, was measured at ages 8 and 16. At age eight it was measured using responses to 12 questions from the shortened version of the Nowicki-Strickland Internal-External (NSIE) scale for preschool and primary children (the non-cartoon format Preschool and Primary Nowicki-Strickland Internal-External scale). At age 16 locus of control was measured using the 12 item Nowicki-Strickland Locus of Control Scale. These scales use questions such as “do you feel that wishing can make good things happen?” and were reverse coded so that high values denote higher locus of control.

#### Empathy

Empathy was measured at age seven using mother reported responses to five questions about the child’s attitudes towards sharing and caring, with responses were on a four-point Likert scale. Responses were coded and summed so that a higher score corresponds to higher levels of empathy.

#### Impulsivity

Impulsivity was measured during two sessions at the age 8 direct assessment using a behaviour checklist. Testers rated whether the children demonstrated restlessness, impulsivity, fleeting attention, and lacking persistence. Responses were coded to 0 (behaviour not characteristic of the child), 1 (behaviour somewhat characteristic of the child), or 2 (behaviour characteristic of the child). Responses were summed and the mean value of the two sessions was used for each child. At age 11 the children were asked a battery of 10 questions designed to capture impulsive behaviour such as “have you spent all of your money as soon as you got it?”. Responses were binary and were summed to give a total impulsivity score.

#### Personality

Personality was measured at age 13 using the five-factor model of personality. Five broad and independent dimensions of personality, the “Big Five” (extraversion, neuroticism, agreeableness, conscientiousness, and intellect), explain a major portion of judged interindividual difference in personality. They are treated as dimensions with individuals varying continuously along these and most people falling between the extreme. They were measured using self-report responses to 50 items of the International Personality Item Pool in which participants indicate the extent to which statements describe their personality on a five-point scale.

### Supplementary Figure 1: Timeline of measures in ALSPAC. SDQ, strengths and difficulties; NEET, not in education, employed or training. See Materials and methods for full details of non-cognitive measures.

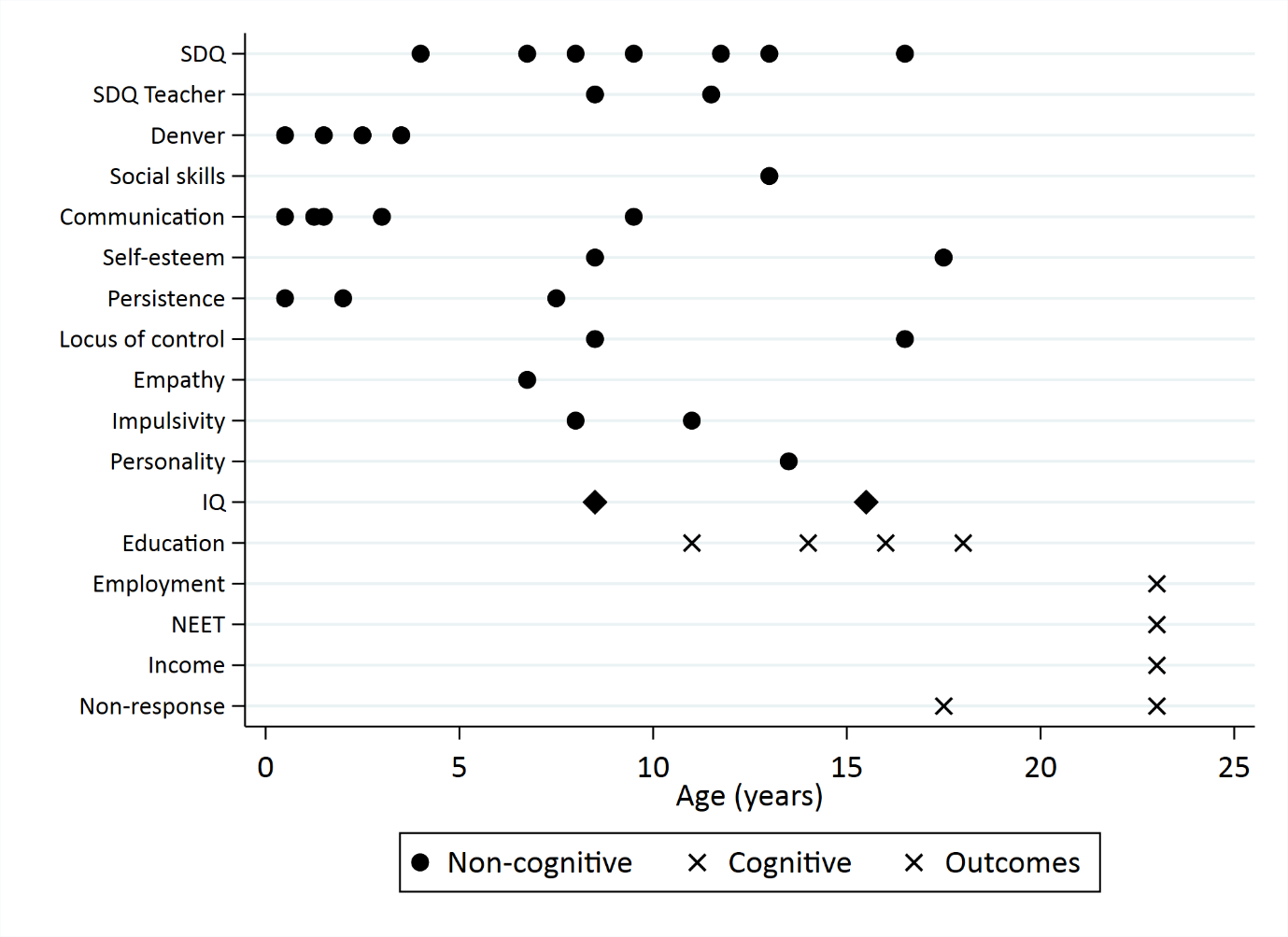

### Supplementary Figure 2: Associations between skills and educational attainment

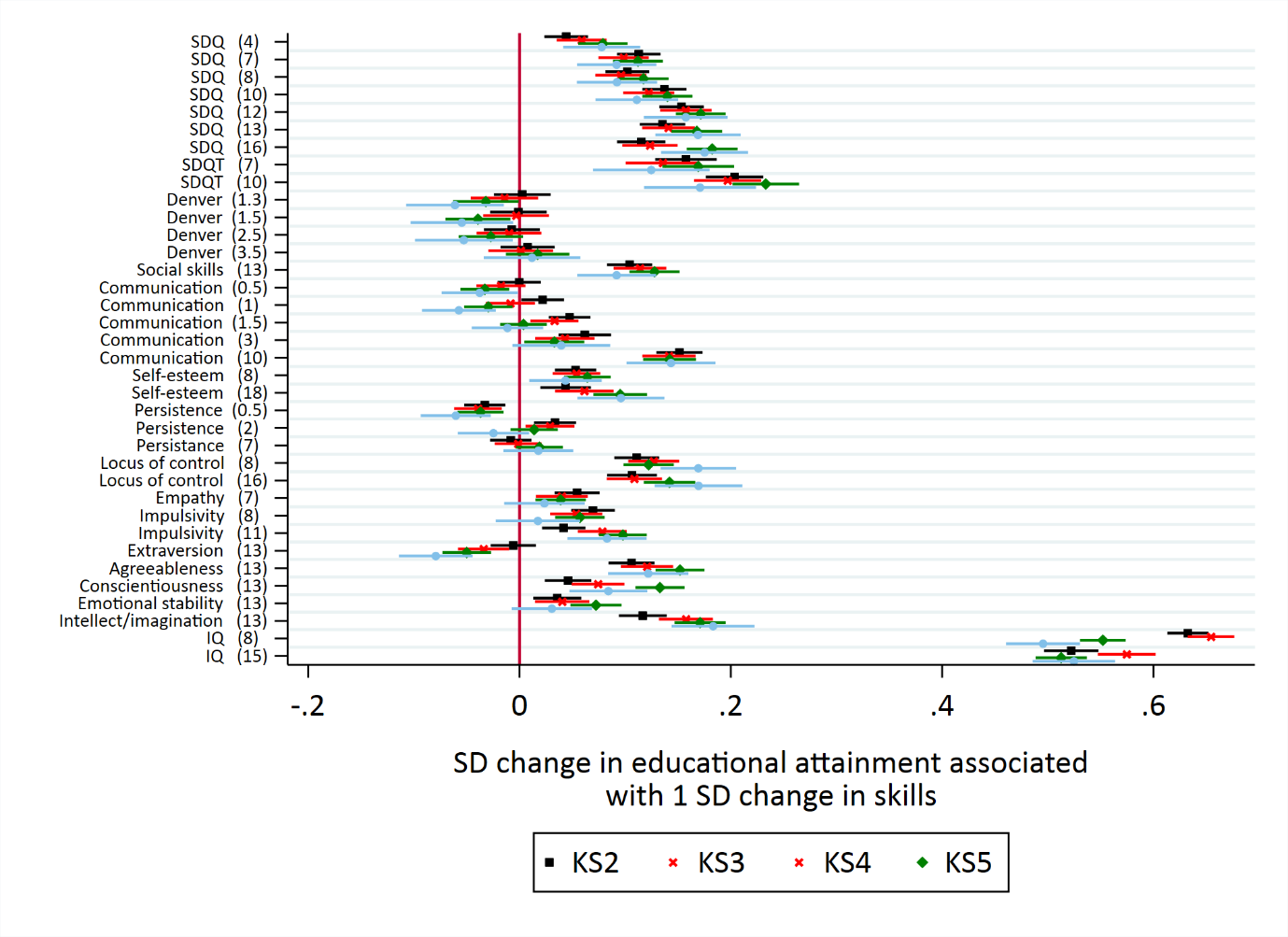

### Supplementary Figure 3: Heritability of SDQ total scores and SDQ internalising/externalising subscales

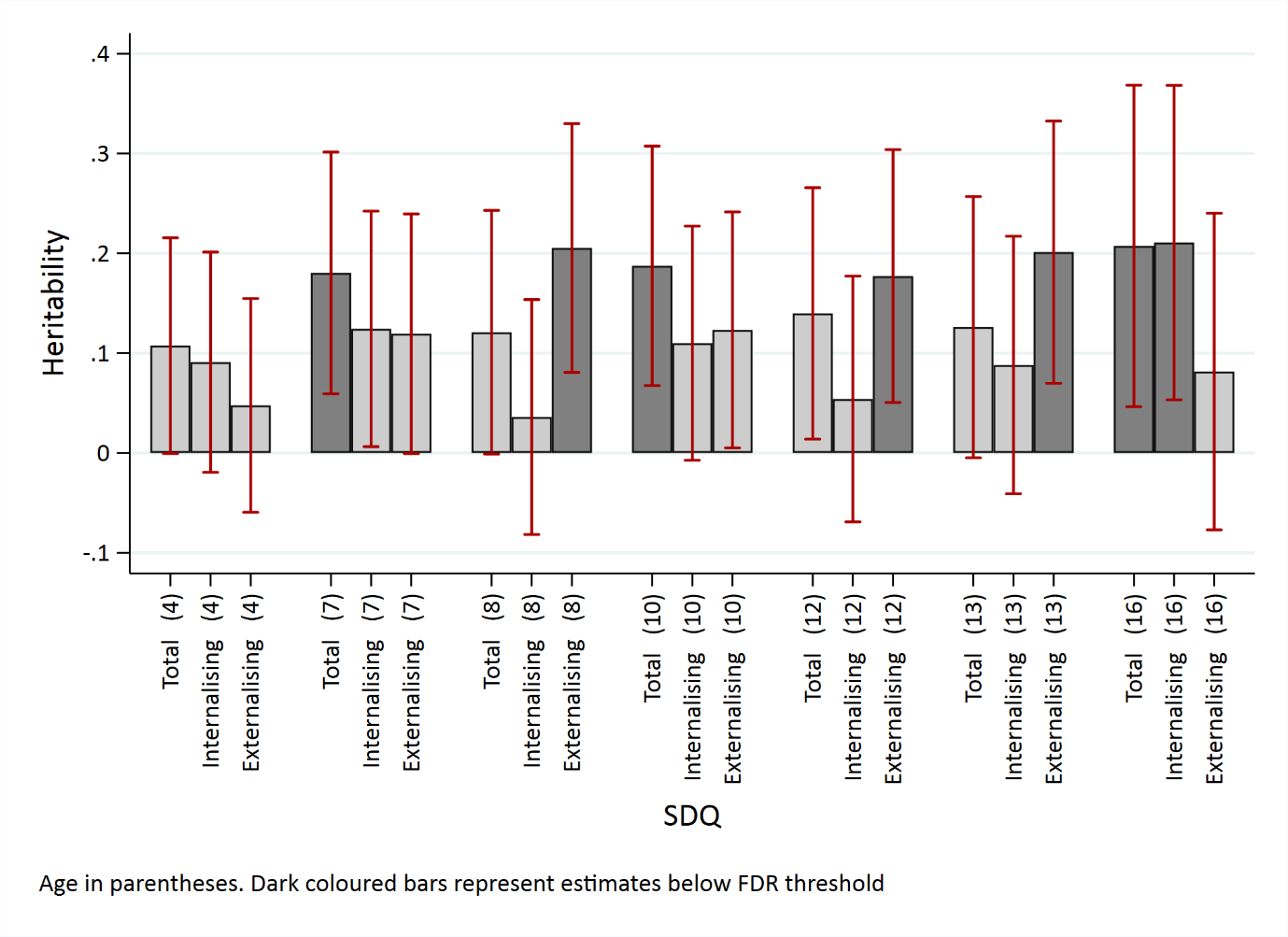
